# Anorectic and anxiogenic actions of cocaine– and amphetamine-regulated transcript in the lateral septum

**DOI:** 10.1101/2024.11.30.626209

**Authors:** Anjali Shankhatheertha, Mikayla A Payant, Jenny Phy-Lim, Melissa J Chee

**Affiliations:** Department of Neuroscience, Carleton University, Ottawa, ON, Canada

**Keywords:** Cocaine– and amphetamine-regulated transcript, melanin-concentrating hormone, lateral septum, feeding, anxiety

## Abstract

Cocaine-and amphetamine-regulated transcript (CART) is produced in several brain regions including the hypothalamus where it is made in cells that also produce melanin-concentrating hormone (MCH). MCH+CART cells densely innervate the lateral septum (LS), which integrates food– and mood-related behaviours. However, while MCH typically promotes feeding and anxiolysis, CART suppresses feeding and promotes anxiogenesis. The LS is a target site of the orexigenic actions of MCH, but it is not known if the actions of CART converge or oppose that of MCH in the LS. We implanted a bilateral cannula over the lateral or central LS of male and female wildtype mice and infused vehicle, CART_55–102_, or co-infused CART and MCH. CART did not alter chow intake but suppressed the intake of a palatable high sugar diet in male and female mice, especially when delivered in the medial LS. Furthermore, CART also prevented the orexigenic effect of MCH on palatable feeding intake when infused in the medial LS. We then assessed if CART regulated anxiety-like behaviour via the LS and found that intra-LS CART infusion reduced time spent in the center of an open field in male but not female mice. Our findings indicated that CART elicited anorectic and anxiogenic actions and may function in opposition to or independently of MCH in the LS. These outcomes suggested that putative CART and MCH co-release from MCH neurons may provide biphasic regulation of feeding and anxiety via the LS.

## 1. Introduction

Neurons that produce cocaine– and amphetamine-regulated transcript (CART) are diverse and widespread. CART-expressing neurons are most prevalent in the ventral regions of the mouse brain,^1^ including the piriform area of the isocortex, amygdalar regions of the cortical subplate, pallidum, and Edinger-Westphal nucleus in the motor-related midbrain, but they are most abundant in the hypothalamus,^2^ especially within the hypothalamic periventricular and lateral zone.^3,4^ One major subset of lateral hypothalamic neurons that express CART produces melanin-concentrating hormone (MCH), as about half of MCH neurons coexpresses CART.^5–8^ MCH/CART+ cells are medially distributed^5,7,8^ and have predominantly anterior projections.^9^ Indeed, the lateral septum (LS) comprises the densest accumulation of projections from MCH cells,^10^ and CART-immunoreactive fibers^11,12^ and varicosities^13^ are distributed throughout the LS rostrocaudally. However, the role of CART in the LS is not well-understood.

MCH and CART are implicated in overlapping physiological roles or behaviours like the control of energy homeostasis, locomotor activity, sleep, and anxiety, but they may have opposing actions on shared roles. For example, MCH and CART are most well-known for their actions on food intake, but while MCH administration can initiate feeding,^14^ prolong food bouts,^15^ or increase food intake per food bout,^16^ central CART administration has powerful anorexigenic actions^17–21^ that can also block the orexigenic actions of Neuropeptide Y.^17^ Furthermore, MCH knockout mice are hyperactive^22^ while CART knockout mice have lower locomotor activity than wildtype littermates.^23^ In sleep-wake regulation, MCH actions promote REM sleep,^24^ while CART promotes wakefulness.^25^ By contrast, both MCH and CART are linked with anxiogenesis. MCH receptor antagonism produces anxiolytic effects^26^ and thus suggests the anxiogenic actions of native MCH.^27^ Meanwhile, CART administration produces a clear anxiogenic response in rodents^28^ that elicits a dose-dependent decrease in the number of open arm or center entries in the elevated plus maze or open field test.^28–30^

The most notable functions of MCH and CART that may overlap at the LS include food-related and anxiety-like behaviours, but it is not clear if MCH and CART signal via the LS to regulate these overlapping functions. The LS is known to integrate food– and mood-related behaviours, thus implying its potential to support the feeding and anxiety effects of CART. LS inhibition elicits orexigenic and anxiolytic outcomes,^31,32^ and we recently found that MCH reversibly inhibits LS cells^33^ and promotes feeding in male and female mice but without affecting anxiety-like behaviours.^34^ As CART can stimulate neuronal activity,^35,36^ we hypothesized that CART may oppose the actions of MCH at the LS to mediate anorectic and anxiogenic outcomes.

We bilaterally infused CART into the lateral or medial LS to resolve putative hotspots for CART action within the LS. CART infusion especially in the medial LS regions suppressed palatable feeding and prevented MCH-mediated feeding in male and female mice. Furthermore, CART infusion also elicited potent anxiogenic actions, which were seen in male mice only. These findings demonstrated that the LS is an important site underlying the anorectic and anxiogenic actions of CART. While it is not known if MCH neurons contribute the main source of CART in the LS, our findings indicated that CART actions in the LS may oppose or be independent of MCH.

## 2. Methods

All procedures were approved by the Carleton University Animal Care Committee according to guidelines outlined by the Canadian Council on Animal Care. All C57BL/6J wildtype mice (stock 000664; Jackson Laboratory, Bar Harbour, ME) were bred and raised in-house on a 12-hour light-dark cycle (22–24 °C) and had *ad libitum* access to water and standard chow (2.9 kcal/g; 2014 Teklad Global Diets, Envigo, Mississauga, Canada).

### 2.1 Cannula implantation

Male and female mice (8–14 weeks old) were given a subcutaneous injection of the analgesic carprofen (20 mg/kg; Zoetis, Kirkland, Canada) at least 30 minutes prior to administering isoflurane anesthesia. Puralube (Dechra, Point-Claire, Canada) eye ointment was applied directly to the eye once the mouse reached anesthetic plane. Fur over the incision site was shaved, and the shaved skin was cleaned with a chlorhexidine scrub and then a 70% isopropyl alcohol wipe. A topical anesthetic or numbing cream (EMLA cream, McKesson Canada, Saint-Laurent, Canada) was applied to the shaved skin for at least 20 minutes. The anesthetized mouse was transferred to a stereotaxic apparatus (David Kopf Instruments, Tujunga, CA) and any excess numbing cream was wiped off the shaved skin. The incision site was sterilized with another round of chlorhexidine, alcohol, and iodine wipe, and a small 2 cm rostrocaudal incision was made to expose the skull.

The skull surface was cleaned with a sterile cotton tip and air-dried. Using one tip of a 26 G bilateral stainless steel guide cannula (C235DCS-5/SPC, Protech International Inc., Boerne, TX) that was fitted with a bilateral mating dummy cannula (C235GS-5/SPC, Protech International Inc.), we read the bregma and lambda coordinates from the top of the skull to level the head position along the horizontal plane within the stereotaxic frame. Additionally, the top of the skull was also leveled along the sagittal plane by reading the coordinates ±1.0 mm to the left and right of bregma. The guide cannula was then moved into position above the desired anterioposterior (AP) and mediolateral (ML) axis relative to bregma, and a 0.5 mm size drill bit was used to drill into the skull at the marked implant location. The stereotaxic coordinates for the medial LS were (in mm): AP +0.7, ML ± 0.4, dorsoventral (DV) −3.3, and the stereotaxic coordinates for the lateral LS were (in mm): AP +0.7, ML ±0.6, DV −3.3. After ensuring that all bleeding had stopped, the guide and dummy cannula complex was slowly lowered to the desired DV coordinate. The cannula was fixed to the top of the skull with two layers of dental cement. Each layer was formed by combining Jet denture repair powder (1230P1, Lang Dental Manufacturing, Wheeling, IL) with Jet self-curing acrylic resin liquid (1404CLR, Lang Dental Manufacturing). The first layer was quickly applied to ensure that at least 2–3 screw turns on the guide cannula were embedded within the dental cement. After ensuring that the first layer was thoroughly dried, we applied a second layer of dental cement so that the guide cannula would be supported by adhering the cement to a larger surface area across the skull. To reduce exposure of the cement base on the skull, the skin around the incision was gently pulled above and sutured around the base of the cannula implant. The dummy and guide cannula were covered and secured with a dust cap (303DC/1, Protech International Inc.) while the mice recovered in their homecage. All mice were closely monitored post-operatively for three days and allowed to recover for at least two weeks before they were habituated to handling for ten consecutive days.

### 2.2 LS infusion

Prior to infusion, the dummy cannula was removed, and a 33 G internal bilateral cannula (C235IS-5/SPC, Protech International Inc.) was inserted inside the guide cannula. Our internal cannula had a 0.25 mm protrusion beyond the tip of the guide cannula and was fitted with thin-walled PE50 tubing (C232CT, Protech International Inc.). The infusate (1 μl total volume per side) was administered over four minutes and allowed another two minutes to diffuse and settle into the target region. The tubing was then removed and switched to the other side for infusion as described. We used a vehicle comprising artificial cerebrospinal fluid (aCSF; in mM: 148 NaCl, 3 KCl, 1.4 CaCl_2_, 0.8 MgCl_2_, 0.8 Na_2_HPO_4_, 0.2 NaH_2_PO_4_). We prepared our infusates, including CART_55–102_ (1 μg; 003-62, Phoenix Pharmaceuticals, Burlingame, CA) or a solution mixture of CART_55–102_ (1 μg) and MCH (1 μg; 4025037, Bachem, Torrence, CA), immediately before use. All infusions started between Zeitberger time (ZT) 1–3.

### 2.3 Food intake

A standard chow or 60% high dextrose diet (3.6 kcal/g; TD.05256 Teklad Custom Diet, Envigo) food pellet was placed in a ceramic soy sauce dish at the bottom of the homecage, and the weight of the remaining food pellet was measured every hour over a 4-h feeding period.

### 2.4 Open field test

Mice were habituated to an opaque, open field box (45 x 45 x 45 cm) at least three times prior to testing. Mice were habituated to the testing room (300 lux) for one hour on test day before starting infusions at ZT1–3. After infusions, mice were returned to their homecage for 30 min and then placed in the corner of the open field arena box, where it may freely explore the entire arena for 10 min. Movement in the open field was tracked using a recording camera (Hue HD camera, Hue, London, UK) placed above the open field, and the camera was connected live to ANY-maze software (Stoelting Co., Wood Dale, IL). The open field was divided into the center zone measuring 22.5 x 22.5 cm and an outer zone, and the mouse was tracked using its center point to determine the duration of time spent in the center or outer zone.

### 2.5 cFos induction

Male and female mice were prepared for stereotaxic surgery, as described in Section 2.1. After drying and leveling the skull, we lowered a glass micropipette to the medial LS using the stereotaxic coordinates (in mm): AP +0.7, ML ±0.5, DV −3.45 and manually injected 100 nl of either aCSF or CART (200 μM) into the separate sides of the LS at a rate of 25 nl/min. We waited at least 5 minutes before retracting the micropipette. Mice remained under anesthesia for 60 min and were euthanized by transcardiac perfusion (see Section 2.6 below).

### 2.6 Tissue processing

Mice aged 8–11 weeks were anesthetized with an intraperitoneal injection of chloral hydrate (700 mg/kg; MilliporeSigma, Burlington, MA) prepared in sterile saline (0.9% NaCl). When mice reached a deep anesthetic plane, they were transcardially perfused with cold (4 °C) sterile saline and then 10% formalin (VWR, Radnor, PA). The brains of the mice were extracted from the skull, post-fixed overnight in 10% formalin (24 h, 4 °C), and cryoprotected in phosphate buffered saline (PBS) containing 20% sucrose and 0.05% sodium azide (24 h, 4 °C). All brains were sliced into five series of 30 µm coronal sections using a freezing microtome (Spencer Lens Co., Buffalo, NY). The sections were immediately mounted onto SuperFrost Plus glass microscope slides (Fisher Scientific, Pittsburgh, PA), and any unmounted series were stored in antifreeze solution^37^ at −20 °C until use.

### 2.7 Indirect immunoperoxidase immunohistochemistry

Free-floating brain sections were washed six times in PBS at RT for 5 min each, treated with 0.3% hydrogen peroxide (H7060-500ML; ACP Chemicals, Montreal, Canada), then washed again three times in PBS for 10 min each. The sections were then blocked in 3% normal donkey serum (NDS; 017-000-121, Jackson ImmunoResearch Laboratories, West Grove, PA; RRID: AB_2337258) for 2 h and immediately incubated with a rabbit anti-c-Fos antibody (1:2,000; ab190289, abcam, Cambridge, UK; RRID:AB_2737414) at RT overnight (16 h). After six 5-min washes in PBS (RT), the slices were incubated at RT for 1 h with a biotinylated goat anti-rabbit antibody (1:500; 111-065-144, Jackson ImmunoResearch Laboratories; RRID:AB_2337965) and then washed again three times in PBS for 10 min each. The sections were then treated with avidin biotin horseradish peroxidase (1:500; PK-6100; Vector Laboratories, Newark, CA) in PBS for 1 h at RT. After another three 10-min washes in PBS (RT), the slices were reacted with 3,3’-diaminobenzidine (SK-4100, Vector Laboratories) and monitored closely for brown colour transformation (<4 min); the reaction was terminated by washing with PBS three times for 10 min each (RT). The stained sections were mounted onto SuperFrost Plus glass microscope slides (Fisher Scientific), air-dried, dehydrated in increasing ethanol concentrations (50%, 70%, 95%, and 100%) for 3 min each, cleared with xylene for 2 h, and coverslipped with Richard-Allan Scientific Mounting Media (Fisher Scientific) using 1.5 thickness glass (22-266-882P; Fisher Scientific).

### 2.8 Brightfield imaging

As previously described,^33^ images of LS-containing brain slices were acquired using a Nikon Ti2-E inverted microscope (Nikon Instruments Inc., Mississauga, Canada) and processed using NIS-Elements Imaging Software (Nikon). Captured images were then exported as TIFF files and imported into Affinity Designer 2 (Serif Europe Ltd, Nottingham, UK) to assess the location of cannula implants or DAB-labeled staining.

### 2.9 Statistical analyses

Mice were assessed using a within-subject design where all infusion conditions were counterbalanced to control for any unexpected sequence effects. Differences in cumulative food intake following aCSF or CART infusion was determined using a repeated measures two-way ANOVA followed by Tukey’s post-hoc testing. Differences in time spent in the center of an open field was compared using a two-tailed paired t test. Sex differences in food intake and open field performance were determined using a three– and two-way ANOVA, respectively. If data points were missing, a mixed-effect model was applied. All descriptive and inferential statistics were determined using Prism 9 (GraphPad Software, Boston, MA), and statistically significant comparisons were determined at *p* < 0.05. Figures were assembled in Affinity Designer 2 (Serif Europe Ltd).

## 3. Results

### 3.1 LS CART infusion had no effect on chow intake in male and female mice

To determine if the LS supports CART-mediated feeding, we bilaterally infused into the lateral LS (**Figure 1A**) of satiated male and female mice and measured chow intake over four hours. All mice increased chow intake over four hours (F(4, 32) = 29.62; p < 0.0001), but compared to aCSF infusion, CART infusion did not alter cumulative chow intake over four hours in male (aCSF: 0.19 ± 0.06 g; CART: 0.18 ± 0.05 g; q = 3.78, p = 0.871; **Figure 1B**) or female mice (aCSF: 0.22 ± 0.05 g; CART: 0.30 ± 0.04; t = 0.98, p = 0.722; **Figure 1C**). There was however a significant main effect of sex (F(1, 31) = 5.16; p = 0.030) and interaction between sex and time (F(4, 31) = 3.36; p = 0.021), as female mice ate more chow than male mice overall.

**Figure 1.**
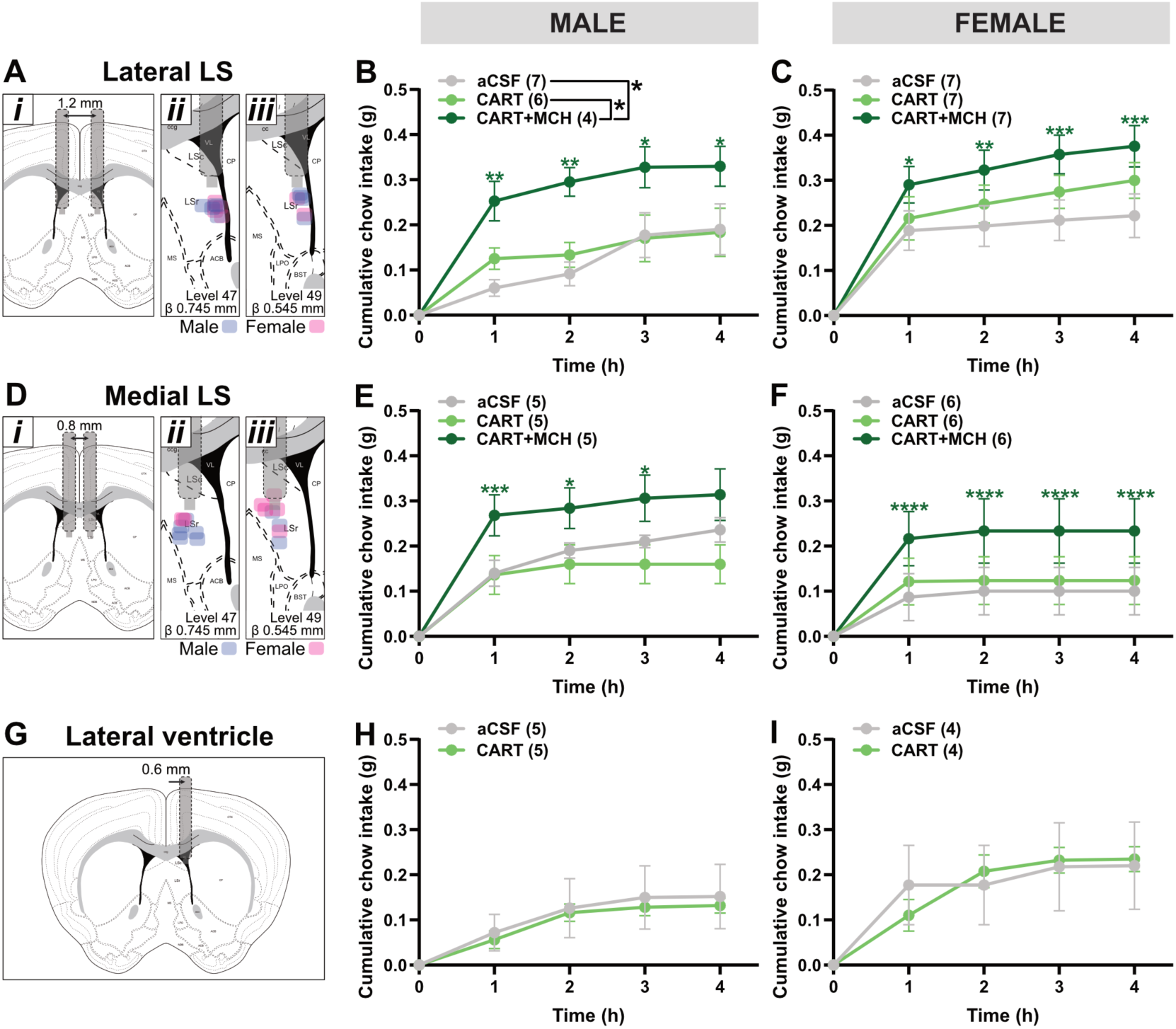
LS CART infusion did not alter chow intake. Schematic of lateral LS infusions with bilateral cannulas implanted within 0.6 mm from the midline on each side (**A*i***). Tip of cannula placement shown on one LS side at *Allen Reference Atlas* level 47 (**A*ii***) and 49 (**A*iii***). Cumulative chow intake following aCSF (1 µl), CART (1 µg/µl), or CART+MCH (1 µg each/µl) infusion into the lateral LS of male (**B**) and female mice (**C**). Schematic of central LS infusions with bilateral cannulas implanted within 0.4 mm from the midline on each side (**D*i***). Tip of cannula placement shown on one LS side at Allen Reference Atlas level 47 (**D*ii***) and 49 (**D*iii***). Cumulative chow intake following aCSF, CART and CART+MCH infusion into the central LS of male (**E**) and female mice (**F**). Schematic of lateral ventricle infusions at the same LS level (**G**). Cumulative chow intake following aCSF and CART infusion into the lateral ventricle of male (**H**) and female mice (**I**). Data are presented as mean ± SEM. Statistical comparisons were made using a repeated measures two-way ANOVA with Tukey’s post-test, where: *, *p* < 0.05; **, *p* < 0.01; ***, *p* < 0.001; ****, *p* < 0.0001 (vs aCSF).

Given that LS MCH infusion increases feeding in male and female mice,^34^ we determined whether CART may oppose or act in synergy with MCH. We bilaterally co-infused CART (1 µg) with MCH (1 µg) into the lateral LS and found a significant increase in chow intake compared to aCSF (q = 4.51, p = 0.015) or CART (q = 3.78, p 0.042) infusion in male mice (**Figure 1B**). The orexigenic effect of CART and MCH co-infusion was not as notable when compared to aCSF (t = 2.37, p = 0.103) or CART (t = 1.39, p = 0.468) infusion in female mice (**Figure 1C**), but there was no sex difference in the orexigenic effect of MCH in the presence of CART (F(1, 33) = 1.56; p = 0.220).

To determine if there may be discrete hotspots for the actions of CART within the LS, we bilaterally infused CART into the medial LS (**Figure 1D**), but CART infusion into the medial LS also did not alter cumulative chow intake in male (aCSF: 0.24 ± 0.03 g; CART: 0.16 ± 0.04 g; q = 0.99, p = 0.771; **Figure 1E**) or female mice (aCSF: 0.10 ± 0.05 g; CART: 0.12 ± 0.05 g; q = 0.65, p = 0.891; **Figure 1F**) compared to aCSF infusion. As seen in the lateral LS, MCH infusion into the medial LS in the presence of CART tended to increase cumulative chow intake relative to CART (MCH+CART: 0.31 ± 0.06 g, n = 6; q = 3.44, p = 0.094) but not aCSF (q = 2.45, p = 0.252) infusion in male mice (**Figure 1E**). In female mice, MCH and CART co-infusion also tended to increase chow intake relative to aCSF (q = 3.29, p = 0.098) and CART infusion (q = 2.64, p = 0.199; **Figure 1F**). The orexigenic actions of MCH in the presence of CART was comparable between male and female mice (F(1, 9) = 1.38; p = 0.270) but was more prominent when delivered to the lateral than medial LS.^34^

We also evaluated the extent of off-target effects from potential drug spillover into the lateral ventricles (**Figure 1G**). However, intracerebroventricular CART infusion did not produce any effects on chow intake over four hours in male (aCSF: 0.15 ± 0.10 g; CART: 0.13 ± 0.10 g; F(1, 8) = 0.07, p = 0.798; **Figure 1H**) or female mice (aCSF: 0.22 ± 0.10 g; CART: 0.24 ± 0.10 g; F(1, 6) = 0.0004, p = 0.985; **Figure 1I**).

### 3.2 LS CART infusion suppressed palatable feeding in male and female mice

While CART did not suppress chow intake, we tested whether the anorexigenic effects of CART would be evident only under hyperphagic conditions. We determined if CART in the LS may suppress hyperphagia induced by access to a palatable 60% dextrose diet^34^ over four hours. Male (chow: 0.17 ± 0.06 g; dextrose: 0.35 ± 0.09 g; p = 0.0006) and female mice (chow: 0.19 ± 0.05 g; dextrose: 0.23 ± 0.08 g; p = 0.009) consumed more dextrose than chow diet over four hours.

Relative to aCSF, lateral LS CART infusion tended to suppress cumulative dextrose intake in male mice (aCSF: 0.77 ± 0.14 g; CART: 0.47 ± 0.12 g; q = 0.97, p = 0.090, **Figure 2A**) but had no effect in female mice (aCSF: 0.68 ± 0.15 g; CART: 0.45 ± 0.09; q = 1.99, p = 0.368, **Figure 2B**). Interestingly, CART co-infusion blocked any orexigenic effect of MCH in both male (CART+MCH: 0.48 ± 0.14 g; q = 3.30, p = 0.080, **Figure 2A**) and female mice (CART+MCH: 0.58 ± 0.13 g; q = 0.478, p = 0.939, **Figure 2B**). There was no sex difference on the anorexigenic effect of CART in the lateral LS on dextrose intake (F(1, 14) = 0.04; p = 0.854).

**Figure 2.**
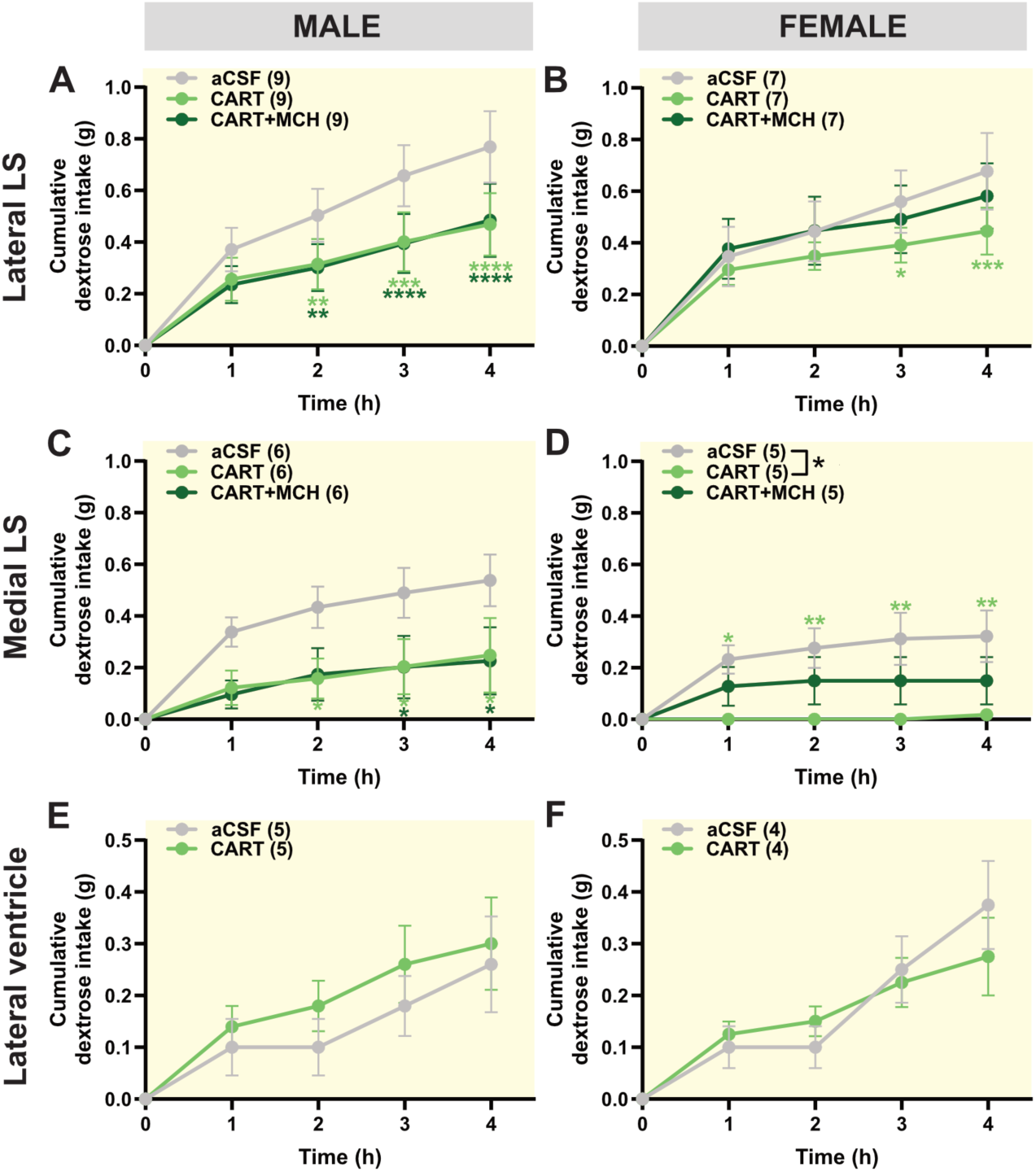
LS CART infusion suppressed palatable feeding. Cumulative intake of 60% high dextrose diet following bilateral aCSF (1 µl), CART (1 µg/µl), and CART+MCH (1 µg each/µl) infusion into the lateral LS of male (**A**) and female mice (**B**) or into the central LS of male (**C**) and female mice (**D**). Cumulative 60% high dextrose diet intake following aCSF and CART infusion into the lateral ventricle of male (**E**) and female mice (**F**). Data are presented as mean ± SEM. Statistical comparisons were made using a repeated measures two-way ANOVA with Tukey’s post-test, where: *, *p* < 0.05; **, *p* < 0.01; ***, *p* < 0.001; ****, *p* < 0.0001 (vs aCSF).

Interestingly, the anorexigenic effect of CART appeared to be more robust in the medial LS. There was a significant main effect of drug in male (F(2, 8) = 5.03, p = 0.039; **Figure 2C**) and female mice (F(2, 8) = 4.36, p = 0.052; **Figure 2D**) that depressed palatable feeding by 66 ± 16% and 92 ± 8%, respectively, relative to aCSF (male: q = 3.82, p = 0.063; female: q = 4.18, p = 0.043). However, the magnitude of the anorexigenic CART effect was not significantly different between male and female mice (F(1, 31) = 3.48; p = 0.072).

CART co-infusion also blocked MCH-mediated palatable feeding. In male mice (**Figure 2C**), CART and MCH co-infusion to the medial LS (0.23 ± 0.13 g) depressed palatable feeding to the same extent as that following CART infusion alone (0.25 ± 0.14 g; q = 0.12, p = 0.996) and was lower than following aCSF infusion (0.54 ± 0.10 g; q = 3.94, p = 0.055). In female mice (**Figure 2D**), MCH infusion in the presence of CART (0.51 ± 0.09 g) also did not elicit palatable feeding relative to that seen following CART (0.02 ± 0.02 g; q = 2.08, p = 0.353) or aCSF infusion (0.32 ± 0.10 g; q = 2.10, p = 0.349).

Feeding following CART and MCH co-infusion was similar between male and female mice (F(1, 31) = 1.26; p = 0.271).

To affirm that anorexigenic CART effects were not attributed to broad central actions due to potential CART leakage into lateral ventricles, we delivered CART directly to the lateral ventricle. Intracerebroventricular CART administration resulted in similar palatable feeding compared to aCSF administration in male (aCSF: 0.26 ± 0.10 g; CART: 0.30 ± 0.10 g; F(1, 8) = 0.55, p = 0.478; **Figure 2E**) and female mice (aCSF: 0.38 ± 0.10 g; CART: 0.28 ± 0.10 g; F(1, 6) = 0.03, p = 0.865, **Figure 2F**) over four hours. Therefore, it is unlikely that the anorexigenic effects of CART occurred at targets away from the LS.

### 3.3 LS CART infusion increased anxiety-like behaviour in male mice

Intracerebroventricular administration of CART can promote anxiety-like behaviour,^28,29^ so we determined whether CART induced anxiogenesis via the LS. As there was no evidence of MCH-mediated anxiety via the LS,^34^ we did not administer CART and MCH co-infusion in our open field test.

CART infusion into the lateral LS decreased time spent in the center of an open field in male mice (F(1, 7) = 6.68; p = 0.036; **Figure 3A**) but not female mice (F(1, 5) = 0.0001; p = 0.992; **Figure 3B**). However, this did not reflect a significant sex difference in open field performance (F(1, 24) = 0.09; p = 0.348). Similarly, CART infusion into the medial LS also lowered time spent in the center of the open field in male (F(1, 2) = 228.6; p = 0.004; **Figure 3C**) but not female mice (F(1, 5) = 0.09; p = 0.777; **Figure 3D**). The difference in open field performance following CART infusion into the medial LS between male and female mice suggested a sex difference in CART-mediated anxiogenesis (F(1, 12) = 3.90; p = 0.072).

**Figure 3.**
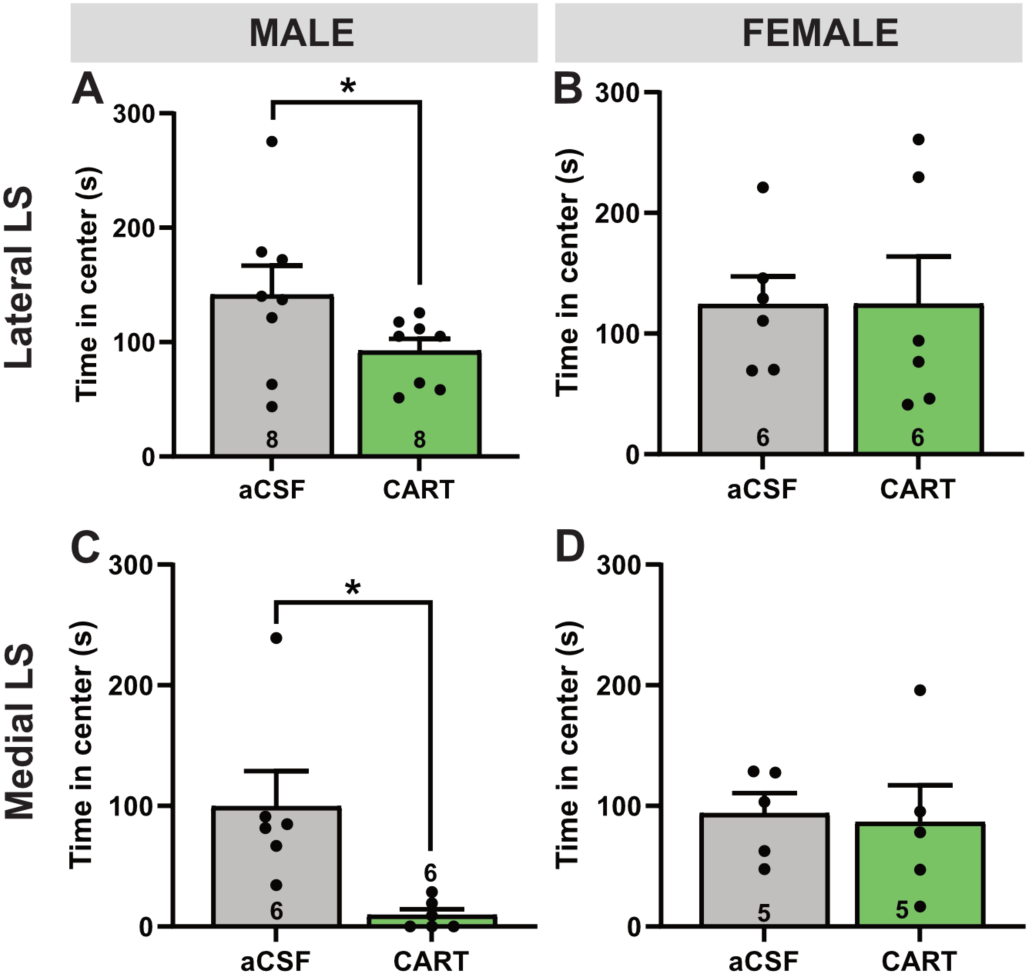
LS CART promoted anxiogenesis in male mice only. Time spent in the center of the open field following bilateral aCSF (1 µl) or CART (1 µg/µl) infusion into the lateral LS of male (**A**) and female mice (**B**) or medial LS of male (**C**) and female mice (**D**). Data are presented as mean ± SEM. Statistical comparisons were made using a paired t test, where: *, *p* < 0.05; **, *p* < 0.01.

### 3.4 CART infusion on LS activity

To determine the functional outcome of CART infusion in the LS on neuronal activation, we unilaterally infused aCSF or CART into the medial LS of the same brain and compared the expression of c-Fos immunoreactivity. There was a greater proportion of c-Fos immunoreactive cells following CART infusion (**Figure 4**).

**Figure 4.**
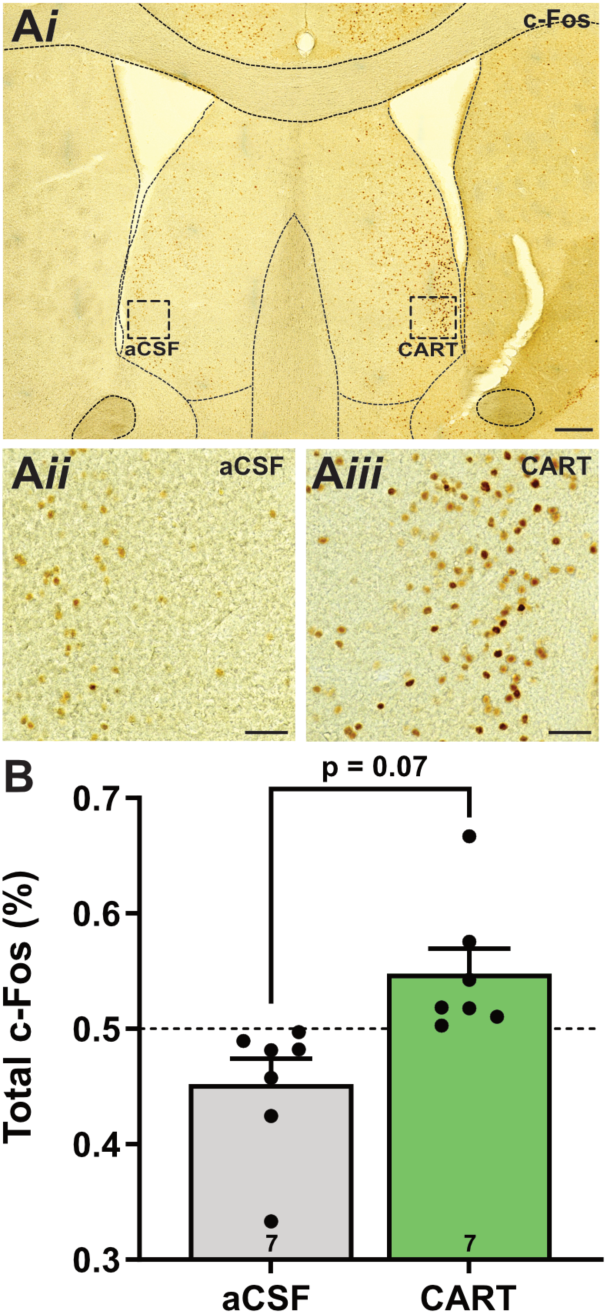
CART infusion increased neuronal activation in the LS. Representative brightfield photomicrograph of c-Fos immunoreactivity following unilateral aCSF and CART infusion into the medial LS of the same brain (**A*i***). High magnification brightfield photomicrographs from dashed outlined region in **A*i*** from the aCSF (**A*ii***) or CART-infused (**A*iii***) LS. Comparison of c-Fos expression expressed as a proportion of total c-Fos immunoreactive cells per brain following aCSF and CART infusion (**B**). Scale bar: 200 µm (**A*i***); 50 µm (**A*ii***, **A*iii***).

## 4. Discussion

This study showed that the LS is a major brain region supporting the anorectic and anxiogenic actions of CART. Bilateral CART infusion directly into the LS suppressed hyperphagia when consuming a highly palatable diet and increased anxiety-like behaviour in the open field test. Interestingly, while the actions of CART on feeding were comparable between male and female mice, the anxiogenic functions of CART were sex-specific, as only CART infusion to male mice elicited anxiogenesis.

The specific source of CART in the LS is not known, but one likely source includes MCH neurons, which have dense LS projections that directly innervate LS cells.^10,33^ Importantly, half of MCH cells coexpress CART.^5,6^ MCH/CART+ cells are medially distributed within the lateral hypothalamus,^5,8^ and medial MCH cells are known to project anteriorly,^9^ such as to the LS. CART-expressing fibers in the LS appear as varicosities,^13^ thus CART may be released extrasynaptically and have actions away from the release site. Nonetheless, CART infusion also elicited a distribution pattern of neuronal activation that included cells along the lateral and ventral regions of the LS, and this activation pattern was consistent with the distribution of both MCH-^33^ and CART-immunoreactive fibers in the LS.^11,12^ Therefore, MCH/CART+ projections may contribute to the availability of CART in the LS.

CART is well-established as an anorexigenic neuropeptide, as intracerebroventricular CART administration acutely suppressed feeding^17–21^ and induced weight loss when administered chronically.^38^ The brain regions associated with the anorexigenic actions of CART include the nucleus accumbens,^39^ hindbrain,^18^ and hypothalamus^40^ though the hypothalamus may also mediate orexigenic actions of CART.^41,42^ In this study, we found that CART suppressed feeding via the LS of male and female mice, but the anorexigenic effect of CART was visible only under hyperphagic conditions, as intracerebral CART administration to the LS suppressed the intake of a highly palatable dextrose diet but not that of standard chow. Furthermore, CART blocked MCH-mediated feeding via the LS,^33^ thus in addition to exerting its own anorexigenic tone, it also suppressed potent orexigenic actions similar to the effect of CART in conjunction with other peptides like Neuropeptide Y.^43^ There were no significant sex difference in the anorexigenic effect of CART in the LS, but the effect of CART tended to be stronger in the female LS, especially in the medial parts of the LS. Previous CART studies used primarily male rodents,^18,20^ so it is not clear if the anorexigenic effect of CART would also be comparable between male and female mice at other sexually dimorphic brain regions. However, our results are consistent with those from CART-knockout mice, where both male and female mice with *Cartpt* gene deletion consumed more calories than littermate controls^44^ thus suggest that food-or metabolism-related processes may be sex-independent.

The anorexigenic effect of CART appeared to be more robust when CART was targeted to the medial region than lateral edge of the LS thus suggesting potential hotspots for CART activity in the LS. While CART fiber immunoreactivity is generally found in the ventral and lateral LS regions,^11,12^ CART may act distal to its release site by volume transmission to its target LS cell, which may be within the medial parts of the LS. As the LS is a periventricular brain structure, we considered that CART may have spilled into the lateral ventricle to mediate anorectic effects via other periventricular brain sites, but we did not observe any evidence for this. Delivering CART directly to the lateral ventricle did not impact chow or palatable food intake. Furthermore, we also considered that our intra-LS CART infusion may have seeped into the nucleus accumbens, but the anorexigenic profile of CART at the LS did not match that of the accumbens. If CART reached the accumbens, we expected that it would suppress chow intake,^39^ but intra-LS CART infusion did not suppress chow intake in our animals. Taken together, our findings support the LS as a novel target site mediating the anorexigenic actions of CART.

In contrast to overlapping functions of CART and MCH in the LS on food intake, MCH had no effect on anxiety,^34^ but CART had a robust anxiogenic effect in the LS. Interestingly, this anxiogenic CART effect was sex-dependent and seen only in male but not female mice. Previous studies administering CART medially have also demonstrated CART-induced anxiogenesis in male rodents only,^28–30^ but similar data in female rodents are lacking. However, the effect of CART on anxiety in female mice can be extrapolated from CART-knockout studies, which also did not observe differences in open field performance in CART-deleted females.^45^ Anxiety is linked with substance use, and CART has been shown to have sex-dependent outcomes in binge drinking where female mice with CART deletion drink less while their male counterparts drink more than their same-sex wildtype littermates^45^; this implicated the CART system as a potential therapeutic target for treating binge drinking in a sex-specific manner.^45^ As the LS is a sexually dimorphic brain region^46–48^ that regulates stress^49–51^ and alcohol intake,^52^ it is a candidate region for sex-specific treatments related to anxiety or alcohol use.

CART increased c-Fos expression in the LS thus suggesting that CART infusion increased neuronal activation in the LS. The specific mechanism underlying the stimulatory actions of CART is not known, but it may result from direct or indirect CART action. First, CART can excite neurons, for example by directly depolarizing the membrane of kisspeptin neurons,^53^ thus CART may directly depolarize LS cells.

Secondly, CART may disinhibit LS cells to increase c-Fos expression^54^ by inhibitory CART signaling via the activation of G_i/o_-protein coupled pathways^55^ and/or inhibition of voltage-gated calcium channels.^56^ The LS comprises primarily GABAergic cells that may form sub-circuits among themselves,^57^ thus CART-mediated inhibition may disinhibit other LS cells.^54^ Importantly, the net action of CART signaling results in LS stimulation, which is associated with anxiogenesis and anorexigenic effects.^31,32^

## 5. Conclusion

The LS is a novel brain region underlying the anorexigenic and anxiogenic actions of CART. In the LS, CART may be released from MCH neurons, and it is interesting that CART and MCH can have opposing effects yet overlapping functions on feeding. MCH inhibits LS cells to drive feeding^34^ while CART has stimulatory actions and suppresses feeding. Our finding that CART can block the orexigenic actions of MCH implicates the potential for biphasic responses in the control of food intake via the LS. As both CART and MCH can have independent or overlapping actions, it would be important for future studies to consider potential interactions due to convergent or divergent outcomes of both peptides at the LS.

## Author Contributions

Conceptualization: MJC. Methodology: AS, MAP, MJC. Investigation: AS, MAP, JPL, MJC. Formal Analysis: AS, MAP, JPL. Visualization: AS. Writing – original draft: AS. Writing – review & editing: AS, MJC

## Acknowledgments

This work was supported by NSERC Undergraduate Student Research Award (AS, JPL), Internship-Carleton University Research Experience for Undergraduate Students (AS, JPL), NSERC Postgraduate Scholarships – Doctoral (MAP), NSERC Discovery Grant (MJC).

